# Fluorescence lifetime imaging of sDarken as a tool for the evaluation of serotonin levels

**DOI:** 10.1101/2024.01.04.574197

**Authors:** Martin Kubitschke, Vanessa Beck, Olivia Andrea Masseck

**Affiliations:** Synthetische Biologie, Univeristät Bremen, 28359 Bremen, Germany

## Abstract

Recent advances in the development of genetically encoded biosensors have resulted in a variety of different neurotransmitter sensors for the precise measurement of the dynamics of neurotransmitters, neuromodulators, peptides and hormones in real time. However, intensity-based measurements of fluorescent biosensors are limited by their dependence on the expression level of the sensor, the intensity of the excitation light, and photobleaching overtime. Here, we show that the FLIM of sDarken (a GPCR-based genetically encoded sensor for serotonin) decreases with increasing serotonin concentrations. Different members of the sDarken family, with different affinities for serotonin, show concentration-dependent changes in fluorescence lifetime according to their dynamic range. We believe that this feature of sDarken is a value-adding complement to intensity-based information and may lead to a better understanding of serotonin dynamics in health and disease.

## Main

Understanding how neural networks generate complex behaviour is one of the great challenges of neuroscience. Neurotransmitters and neuromodulators dynamics modulate neuronal circuits, and understanding their function is key to unravelling their role in behaviour. Many diseases are associated with dysfunctional neuromodulatory signaling. Especially the neuromodulator serotonin (5-HT) plays an important role in a variety of physiological functions, cognition and behavior. 5-HT seems to be involved in regulating emotional and social behaviors and disturbances in brains serotonin levels are associated with psychiatric disorders, such as major depressive disorder (MDD). Although the function of 5-HT has been studied for many years, the lack of appropriate tools to measure dynamics on different timescale with high spatial resolution prevented the detailed elucidation of underlying disease mechanisms. A special feature of neuromodulators, such as serotonin (5-HT), is that they act not only as classical neurotransmitters at synapses on a millisecond timescale, but also through volume transmission on the scale of seconds to minutes. These longer timescale dynamics of serotonin are thought to underlie internal brain states or behavioral states, such as sleep, foraging or escape ^1-3^.

In recent years, an increasing number of genetically encoded fluorescent biosensors have been published that allow the measurement of neurotransmitter release with high temporal and spatial resolution in vitro and in vivo ^4-7^. All of this intensity or intensiometric biosensors have in common that upon binding of the ligand to the sensing moiety, conformational changes occur that can either quench or enhance fluorescence. A change in fluorescence can be mediated by several factors, such as shifting the protonation equilibrium of the chromophore, changing the quantum yield or the extinction coefficient ^8^.

However, intensity-based sensors all suffer from several limitations, such as their dependence on expression levels, excitation power, photobleaching and sensitivity to pH changes^9^, which will limit the possible interpretations that can be derived from the intensity readout. A more robust readout may be fluorescence lifetime information (FLIM), which is not affected by expression level, excitation power, photobleaching or pH ^10,11^. Only tools that will enable the precise measurement of serotonin dynamics independent from expression levels of the sensor and excitation power will make it possible to compare serotonin levels across animals, brain areas, disease states and timepoints.

Previously it has been shown that some, but not all GPCR based sensors for neuromodulators, exhibit changes in their lifetime upon binding of their respective ligand and can be utilized to compare neuromodulator dynamics across animals, brain regions, disease models and over time ^12^. Although Ma et al. reported that also GRAB-5-HT showed a change in lifetime upon saturating serotonin concentrations, detected changes were quite small (<0.1 ns), hindering the ability of GRAB5-_HT_ to be utilized as FLIM based sensor.

We hypothesized that sDarken, a GPCR-based intensiometric biosensor that decreases its fluorescence upon binding of serotonin, would also show changes in fluorescence lifetime upon application of 5-HT (**Fig. 1a**). To test our hypothesis, we applied increasing concentrations of serotonin to human embryonic kidney (HEK) cells transiently expressing sDarken.

**Figure 1:**
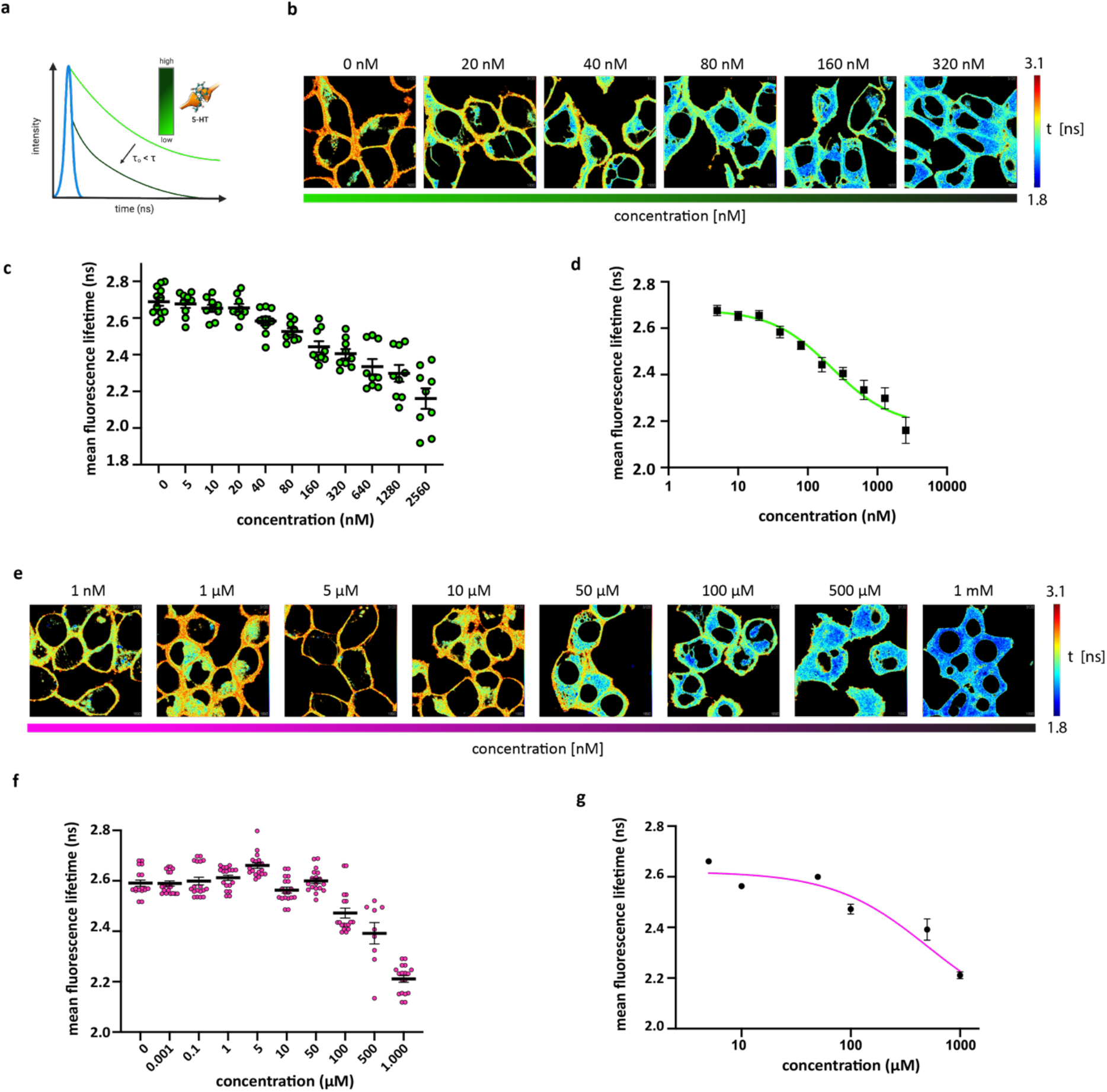
Fluorescence lifetime decreases with increasing concentration of serotonin. (a) Schematic showing the proposed change in fluorescence lifetime by increasing concentrations of serotonin. (b) Representative heatmaps of fluorescence lifetime of sDarken in response to increasing concentrations of serotonin. (c) Mean fluorescence lifetime in response to increasing concentrations of serotonin. 0 nM: 2.689 ± 0.0022 ns, N = 721; 5 nM: 2.676±0.022 ns, N = 810; 10 nM: 2.653±0.019 ns, N =854; 20 nM: 2.655±0.021 ns, N=677; 40 nM: 25584 ±0.024 ns, N=467; 80 nM: 2.526 ±0.017 ns, N = 702; 160 nM: 2.443± 0.030ns, N= 744; 320 nM: 2.405±0.025 ns, N =399; 640 nM: 2.335±0.041 ns, N= 566; 1280 nM: 2.298±0.045ns, N= 618, 2560nM: 2.160±0.055, N=632 9replicates for each concentration, linear x-axis. (d) Inhibitory dose response curve, logarithmic x-scale. (e) Representative heatmaps of fluorescence lifetime of L-sDarken in response to increasing concentrations of serotonin. (f) Mean fluorescence lifetime of L-sDarken in response to increasing concentrations of serotonin. 1 nM: 2.680 ±0.0022 ns, N=154;; 10 nM: 2.684±0.0022 ns, N=131; 100 nM: 2.2.681±0.0022 ns, N= 142; 1 μM: 2.674±0.0021 ns, N=171;5 μM: 22.731±0.005 ns, N=81; 10 μM: 2.659±0.0025 ns, N= 123; 50 μM: 2.684±0.0011 ns, N=124; 100 μM: 2.581±0.0018 ns, N=220; 500 μM: 2.532±0.0032, N=89; 1mM: 2.218±0.0047 ns, N=113) Linear x-axis. n=9 replicates. (g) Inhibitory dose response curve, logarithmic x-scale. (g). mean ± S.E.M.

We observed that the fluorescence lifetime is decreased with increasing concentrations of 5-HT (**Fig. 1b-c**). Lifetime was significantly different from baseline starting at a concentration of 80 nM (p< 0.0021, One Way ANOVA). It is noteworthy that changes in FLIM are only observed within the dynamic range of sDarken (Kd = 127 nM; Kubitschke et al. 2023). The dynamic range of sDarken, as characterized in Kubitschke et al., is between 100 nM and 1 μM, which corresponds well to a decrease of FLIM within this concentration range. We observed that the dynamic range (0.529 ns) of the lifetime between baseline (τ=2.689± 0.0022 ns)and the maximum concentration of 2.560 μM (τ=2.160±0.055 ns) is relatively large. The dose-response curve displays a sigmoidal shape (**Fig. 1d**) and an IC50 of 221.4 nM.

To verify our hypothesis, that changes in FLIM will only be observable with the dynamic range of the utilized biosensor we repeated our experiments with L-sDarken a low affinity variant with 1.000 fold lower affinity for serotonin (KD =45 μM, Kubitschke et al. 2022). In deed changes of the lifetime could only be observed between 10 μM and 1 mM (Fig.1 e-g), well in accordance with the reported dynamic range of L-sDarken. Fluorescence lifetime is decreased with increasing concentrations of 5-HT (**Fig. 1e-f**). Lifetime was significantly different from baseline starting at a concentration of 50 μM (p< 0.021, One Way ANOVA). As for sDarken, also L-sDarken had a remarkable large dynamic range of 0.462 ns, between the baseline lifetime (mean τ=2.680 ±0.0022 ns) and a maximum concentration of 1 mM (mean τ=2.218±0.0047 ns). For control we used a nullmutant of sDarken and a cytosolic expressed GFP. The nullmutant of L-sDarken combines several mutations in the binding pocket of the 5-HT_1A_ receptor the sensing domain of sDarken: A50V, D82N, D116N, S198A and T199A, that have been shown to be important for the binding and activation of the 5-HT_1A_ receptor ^13,14^. Both controls did not show any significant changes in their lifetime to different concentrations of serotonin (**Fig.2 a-b**).

**Figure 2:**
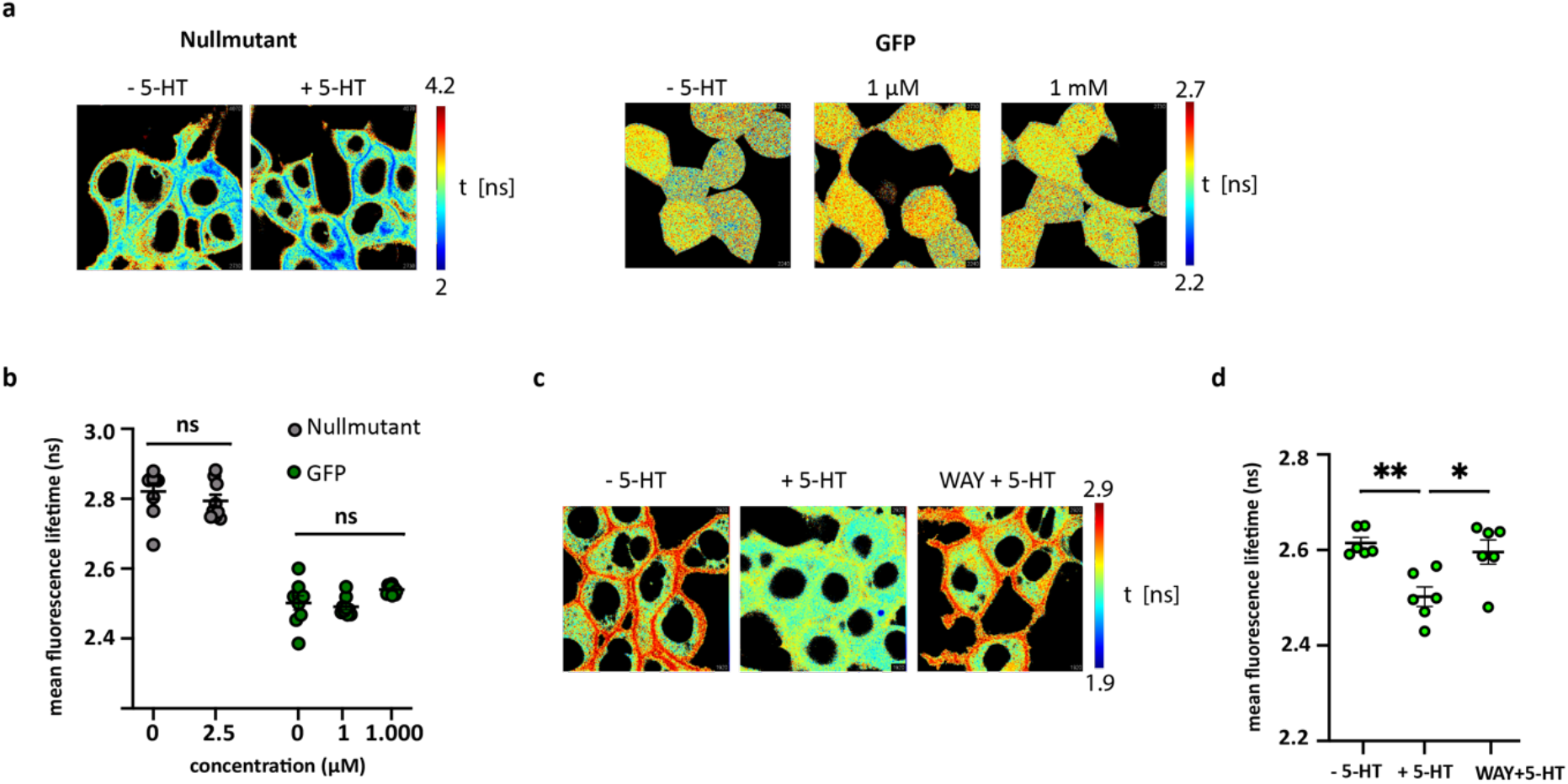
Controls showed no changes in FLIM to 5-HT application. (a) Representative heatmaps of fluorescence lifetime of a sDarken nullmutant and cytosolic expressed GFP in response to saturating concentrations of serotonin (b) Mean fluorescence lifetime of controls in response to different concentrations of serotonin. -5-HT nullmutant: mean τ=2.669± 0.0022 ns, n=107; 2560 nM nullmutant: mean τ=2.743± 0.0018 ns n=85; - 5-HT GFP: mean τ=2.386± 0.0020 ns n=25 ; 1 μM GFP: mean τ=2.469± 0.0008 ns, n=31; 1mM GFP mean τ=2.540± 0.0004 ns, n= 22; 9 replicates (c) Representative heatmaps of fluorescence lifetime of sDarken in response to serotonin (+ 5-HT) and application of an 5-HT_1A_ receptor agonist WAY. (d) Mean fluorescence lifetime of sDarken in response to 10 μM 5-HT and 10 μM WA. – 5-HT: mean τ=2.615± 0.0011 ns, N =458; + 10μM 5-HT : mean τ=2.430± 0.0020 ns, N = 417; 10 μM WAY +10 μM 5-HT : mean τ=2.596± 0.00251 N= 408; 6 replicates. mean ± S.E.M.

The decrease in lifetime is blocked by the application of the 5-HT_1A_ antagonist WAY (**Fig.2 c-d**). Lifetime is significantly decreased with the application of 5-HT (−5-HT: mean τ=2.615±0.0011 ns; + 5-HT: mean τ=2.502±0.0020 ns; one-way ANOVA p=0.003), whereas in the presence of WAY no change in lifetime could be observed (−5-HT: mean τ=2.615±0.0011 ns; WAY+5-HT: mean τ=2.596 ±0.00125 ns; one-way ANOVA p=0.7832).

In conclusion, sDarken, a genetically encoded serotonin sensor, exhibits a concentration-dependent change in fluorescence lifetime. To date, sDarken has the largest reported lifetime change for GPCR based neuromodulator sensors. For sDarken lifetime changes are observed at submicromolar 5-HT concentrations and across micromolar concentrations for L-sDarken, the low affinity variant of sDarken. We suggest that in the future, FLIM measurements of the sDarken family could be a valuable complement to intensity-based measurements of serotonin. As been shown for GRAB_ACh3.0_ ^15^, sDarken could provide additional information about tonic levels of serotonin during different internal states ^12^. FLIM measurements of sDarken have the potential to quantitatively assess serotonin levels, also over long periods of time making it an ideal tool to investigate differences in brain serotonin levels in health and disease. Another advantage would be that FLIM as a readout could presumably be used to make better comparisons of serotonin levels across brain regions, animals, and disease states^12^.

## Acknowledgements

We would like to thank Andreas Reiner and Tommaso Patriarchi for valuable discussions. We thank Celina Schreiber for expert technical assistance. O.A.M and M.K. were funded by the Deutsche Forschungsgemeinschaft (DFG MA 4692/6-3).

## Declaration of generative AI and AI-assisted technologies in the writing process

During the preparation of this work the author(s) used DeepLWrite in order to improve language and readability. After using this tool/service, the author(s) reviewed and edited the content as needed and take full responsibility for the content of the publication.

## Material and Methods

### Cell culture

HEK cells (HEK293, tsA201 cells, ATTC) were cultured in Dulbecco’s modified Eagle’s medium (DMEM, high glucose, with stable glutamine, with pyruvate, Cellpure®) supplemented with penicillin (100 I.U/mL, Cellpure®), streptomycin (0.1 mg/mL, Cellpure®) and fetal bovine serum (Brazilian origin, Gibco). Cells were seeded on μ-Dishes (35mm μ-Dish, IbiTreat, Ibidi) and transfected when they reached ∼70% confluence. For transfection, 1μg plasmid DNA and 4μL polyethyleneimine (branched, average MW 25000, Aldrich Chemistry) were incubated in 100μL DMEM for 15min at RT. The mixture was applied dropwise to 60-70% confluent HEK cells. The cells were incubated at 37°C with 5% CO2. Measurements were performed the following day.

#### Plasmids

Following plasmids were used for transfection pN1-CMV-sDarken (Addgene plasmid #184799, pN1-CMV-L-sDarken (Addgene plasmid #184800), null-mutant of sDarken (created in house), peGFP-N1-FLAG (Addgene plasmid # 60360).

### Measurement

Serotonin stock solution (serotonin hydrochloride, 100 mM in PBS [0.9 mM CaCl2, 0.5 mM MgCl2, 2.7 mM KCl, 1.47 mM KH2PO4, 138 mM NaCl, 8 mM Na2HPO4]) was prepared fresh each day before measurement. Different concentrations of serotonin were prepared by diluting the stock solution with 1X PBS. For each dish, the medium was aspirated and the dish was washed with 1x PBS. Then 2 mL of serotonin solution was added to the cells. For WAY-100635 measurements, 2 μL of solution (10 μM; serotonin, 10 μM WAY-100635 [WAY-100635 Maleate, Sigma Aldrich]) was pipetted directly onto the cells between lifetime measurements. Cells were measured under a confocal laser scanning microscope (Microscope: LSM880 [Zeiss] with the LSM upgrade kit [Picoquant], emission filter 520/35). The measurement was performed with a pixel dwell time of 8.19μs and a frame size of 512x512 pixels. Excitation was performed with a 480nm laser (LDH-D-C-485, Picoquant) at a pulse rate of 40.00MHz. The laser intensity was set just below 10% of the laser repetition rate. Measurements were stopped at 1000 counts in the brightest pixel. The laser power was attenuated to remain below 10% of the laser repetition rate (4000 counts/s).

### IRF measurement

The instrument response function was measured with a saturated NaI solution containing fluorescein (Thermo Fischer Scientific) using the same measurement parameters as above.

### Analysis

FLIM fitting was performed using the FLIMfit analysis software (FLIMfit 5.1.1). Images were loaded with a time binning of 250. The IRF shift was determined by an initial fit and subtracted for fitting. Fitting was performed on thresholded images, and the raw mean tau values of the images were used to determine the mean lifetime of fluorescence within outer membrane organelles. N were selected using the analyze particle function of ImageJ (ImageJ 1.53c). The threshold was set to include most of the membrane parts. N of non-membrane parts were manually discarded.

### Statistics

All statistical analysis was done in GraphPadPrism9. A one way ANOVA was utilized to test whether or not the means of independent sample groups are different. One-way ANOVA assumes normality and homoscedasticity of the data sets. The normality of the data was confirmed by the Shapiro-Wilk test. Homogeneity of variances between groups was tested using Bartlett’s test. Dose response curves are fitted with a dose-response-inhibition (three parameters) fit. Inhibitor vs. response. A standard slope is considered equal to a Hill slope with a sigmoidal shape.

